# Cohesin complex oligomerization maintains end-tethering at DNA double-strand breaks

**DOI:** 10.1101/2023.11.08.566226

**Authors:** Jamie Phipps, Mathias Toulouze, Cécile Ducrot, Rafaël Costa, Clémentine Brocas, Karine Dubrana

## Abstract

DNA double-strand breaks (DSB) must be repaired to ensure genome stability. Crucially, DSB ends must be kept together for timely repair. In *Saccharomyces cerevisiae*, two poorly understood pathways mediate DSB end-tethering. One employs the Mre11-Rad50-Xrs2 (MRX) complex to physically bridge DSB ends. Another requires the conversion of DSB ends into single-strand DNA (ssDNA) by Exo1, but the bridging proteins are unknown. We uncover that cohesin, its loader and Smc5/6 act with Exo1 to tether DSB ends. Remarkably, cohesin specifically impaired in oligomerization fails to tether DSB ends, revealing a new function for cohesin oligomerization. In addition to the known importance of sister chromatid cohesion, microscopy-based microfluidic experiments unveil a new role for cohesin in repair by ensuring DSB end-tethering. Altogether, our findings demonstrate that oligomerization of cohesin prevents DSB end separation and promotes DSB repair, revealing a novel mode of action and role for cohesin in safeguarding genome integrity.

## Introduction

DNA double strand breaks (DSBs) pose a significant threat to genome stability as they disrupt chromosome integrity. Repair mechanisms, such as NHEJ and homologous recombination are essential for restoring chromosome continuity by directly rejoining DSB ends or using a donor homologous template (*1*). However, before these repair processes can occur, it is imperative to bring DSB ends together, a task unlikely achieved through passive diffusion (*2*). Instead, active DSB end-tethering mechanisms have been identified, and represent a critical step in preventing joining or recombination events between unrelated chromosome loci, which could lead to harmful translocations.

The mechanisms that facilitate the tethering of DSB ends were initially characterized in the budding yeast *Saccharomyces cerevisiae* (*3*–*5*). The MRX^MRN^ complex (<u>M</u>re11-<u>R</u>ad50- <u>X</u>rs2^NBS1^) is rapidly recruited to DSB ends and plays an early role in end-tethering (*3, 5, 6*). MRN has been proposed to serve a similar tethering function, thus preventing translocations in humans (*7*–*9*). In yeast, MRX nuclease activity is dispensable for DSB end-tethering. Instead, the ZN-hook domain and ATPase activity of Rad50 are essential, suggesting a physical bridging mechanism by MRX dimers (*3*). In contrast, during later stages of repair, DSB end-tethering requires Exo1 exonuclease activity to reveal single stranded DNA (ssDNA) (*4*). However, the proteins responsible for physical bridging of DSB ends during these late stages of repair remain unidentified.

Recent theoretical research has proposed a role for DNA loop extrusion in the tethering of DSB ends (*2*). Loop extrusion, a property associated with SMC family complexes, has emerged as a conserved mechanism for folding the genome (*10*). Among these SMC complexes, cohesin (comprising Smc1, Smc3, Mcd1^Scc1^ and Scc3^STAG1/2^) and the Smc5/6 complex are recruited to DNA damage sites in both yeast and mammals (*11*–*14*). In yeast, the loading of cohesin to DSBs involves various factors, including the cohesin loader Scc2/Scc4, DNA damage factors like MRX, γH2A, Tel1^ATM^, Mec1^ATR^, and Smc5/6 (*13*–*17*). Cohesin is not only enriched at DSB sites, but also throughout the entire genome (*18*–*20*), contributing to tightening of sister chromatid cohesion (*18, 19, 21*–*23*), locally restricting homology search (*24*), and aiding in DNA damage checkpoint establishment (*20*).

Given the involvement of cohesin in DSB response (*25*), its demonstrated ability to bridge DNA molecules *in vitro* (*26*), and the observed gross chromosomal rearrangements and translocations in cohesin mutants (*27*), we hypothesized that cohesin and/or Smc5/6 play a critical role in DSB repair by maintaining proximity between DSB ends.

In this study, we combine genetic and live microscopy-based approaches to demonstrate a cohesin dependent DSB end-tethering mechanism, involving Exo1 and Smc5/6. Furthermore, we show that cohesin compacts DSB adjacent chromatin, beyond compaction observed in G2/M cells. We expose oligomerization as a key mechanism for both MRX- and cohesin-dependent tethering through both disruption of protein-protein interactions in response to hexanediol treatment, and genetic loss of function mutants. Specifically, disruption of cohesin oligomerization through mutation in the Mcd1^SCC1^ subunit, maintains compaction at the vicinity of DSB, but prevents the ability to tether DSB ends. Disruption of oligomerization between Rad50 heads also leads to loss of MRX dependent DSB end-tethering. Finally, our real-time microfluidic assay demonstrates that cohesin is essential for efficient repair of DSBs, through its end-tethering capacity.

## Results

### Cohesin Tethers DSB ends

To assess the requirement of cohesin in tethering DSB ends, we developed a microscopy-based assay in which LacO and TetO repeats were positioned either side of the endogenous HO endonuclease cleavage site at the *MAT* locus of *Saccharomyces cerevisiae* (Fig. 1A; (*28*)). Targeted by LacI-mCherry and TetR-GFP fusion proteins, these arrays allow for visualization of the regions flanking the DSB site as red and green spots. DSBs were induced by galactose treatment, which triggers the Gal promoter-controlled expression of the HO endonuclease (fig. S1A). In individual cells, we distinguished tethering or separation of DSB ends based on the distance between the spot centers being less than or greater than 400 nm (Fig. 1B). This threshold was established by quantifying spot separation in the absence of DNA DSB, where less than 5% of WT cells exhibited spots exceeding 400nm separation (fig. S1B). We confirmed the assay’s sensitivity to detect the previously described, early MRX-, and late Exo1-dependent end-tethering pathways by imaging at 2 hours and 4 hours post-DSB induction. At 2 hours post-DSB, WT and *exo1Δ* cells showed less than 10% untethering, while cells lacking Mre11 displayed 31% separation (Fig. 1C). At 4 hours post-DSB, separation remained unchanged in WT cells but increased to 23% in *exo1Δ* cells. Importantly, double deletion of *EXO1* and *MRE11* led to a significant increase in end separation compared to either single mutation, highlighting the presence of two pathways of DSB end-tethering (Fig. 1D).

**Fig. 1.**
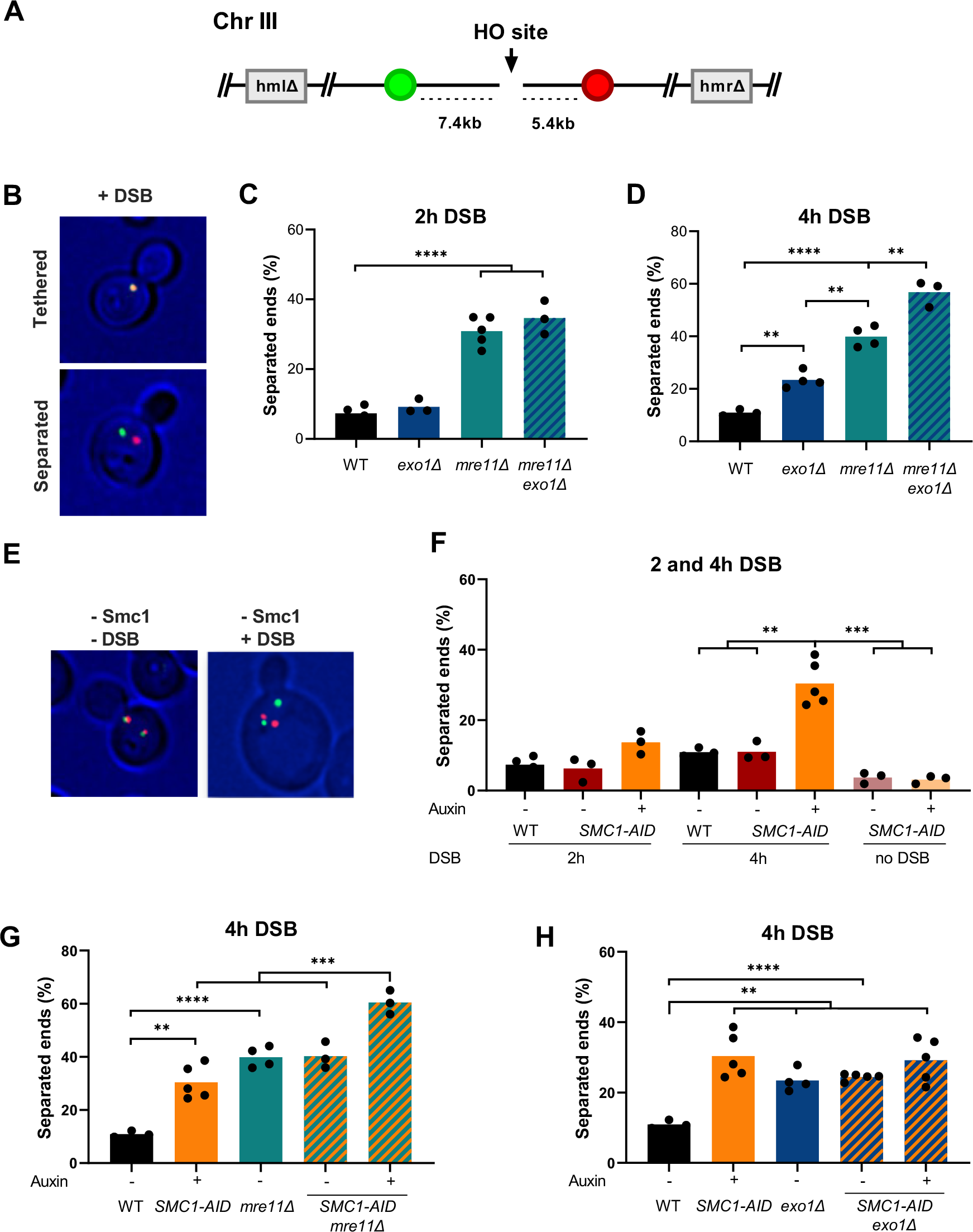
Cohesin Tethers DSB ends in the Exo1 pathway. **(A)** LacO/LacI-mCherry tag and a TetO/TetR-GFP tag were inserted at 5 and 7 kb from the HO DSB site at the MAT locus respectively. (B) Example of cells with tethered or separated ends. Signals are considered as separated when the distance between centers is more than 400nm. (C-D) Percentage of cells with separated ends in the indicated strains after 2 hours (B) or 4 hours (D) DSB induction. (E) Examples of cells showing sister chromatid separation and DSB end separation upon Smc1- AID auxin mediated degradation in absence or in presence of DSB induction. (F) Percentage of cells with separated ends in WT and SMC1-AID strains in absence (-) or presence (+) of auxin after 2 hours, 4 hours or no DSB induction as indicated. (G-H) Percentage of cells with separated ends in the indicated strains after 4h DSB induction. Black stars indicate statistical differences (* = p<0,05; ** = p<0,01; *** = p<0,005; **** = p<0,001).

To investigate the role of cohesin in DSB end-tethering, we employed the auxin-induced degron (AID) system to deplete the cohesin subunit Smc1 (*29*). Following a 1-hour auxin incubation, Smc1 protein levels were substantially reduced and maintained at near undetectable levels for over 4 hours (fig. S2A). Depletion of Smc1 resulted in the appearance of cells with separated sister chromatids (Fig. 1E) and impaired cell growth (fig. S2E), consistent with the essential role of the cohesin complex in sister chromatid cohesion. At 2 hours post-DSB, a non-significant increase in end separation was observed upon cohesin depletion (Fig. 1E). However, at 4 hours post-DSB, approximately 30% of DSB ends were untethered (Fig. 1F). To ensure that the increase in spot separation above 400 nm was due to the lack of DSB end-tethering and not due to the loss of cohesin-mediated chromatin folding, we quantified the percentage of cells with spots exceeding 400 nm upon Smc1 depletion in the absence of DSB. No significant increase in spot separation was observed when Smc1 was depleted in absence of DSB (Fig. 1F), excluding an involvement of cohesin-mediated chromatin folding. Overall, these results reveal the requirement of cohesin for DSB end-tethering.

### Cohesin tethers DSB ends in the Exo1 pathway

To determine the specific pathway in which cohesin tethers DSB ends, we quantified the extent of DSB end separation upon depletion of cohesin in cells lacking Mre11 and Exo1. In contrast to depletion of Smc1 alone, loss of Smc1 in *mre11Δ* cells significantly increased end separation at 2 hours post-DSB (fig. S1C). This early Smc1-dependent end-tethering separation is not seen in *mre11Δ exo1Δ* double mutants, or *exo1Δ* cells depleted for Smc1 (fig. S1D), suggesting that, at 2 hours, cohesin acts in parallel to MRX, independently of Exo1, to maintain end-tethering. Strikingly, at 4 hours post-DSB, depleting cohesin in *mre11Δ* cells significantly increased end separation (Fig. 1G), recapitulating the separation observed in *mre11Δ exo1Δ* cells (Fig. 1D). In contrast, depletion of Smc1 in *exo1Δ* cells did not further increase end separation compared to *exo1Δ* cells (Fig. 1H). These findings suggest that cohesin functions with Exo1 to tether DSB ends at 4 hours post-DSB.

### Cohesin DSB end-tethering requires *de novo* cohesin loading

Next, we wondered if the presence of cohesin on chromosomes prior to DSB induction is sufficient for maintaining DSB end-tethering, or if the DSB-induced *de novo* loaded population of cohesin is required for this function. To address this, we arrested cells in G2/M phase using nocodazole, depleted Scc2 for 1 hour to prevent *de novo* cohesin loading while maintaining pre-existing loops (*30*) and induced DSB (Fig. 2A, and fig. S2B). We observed an increase in separated ends after 4 hours DSB induction upon Scc2 depletion (Fig. 2B), reaching a similar level as that observed in Smc1-depleted and *exo1Δ* cells under the same experimental settings. These results indicate that preformed cohesin loops are not sufficient and that *de novo* loading of cohesin is necessary for DSB end-tethering.

**Fig. 2.**
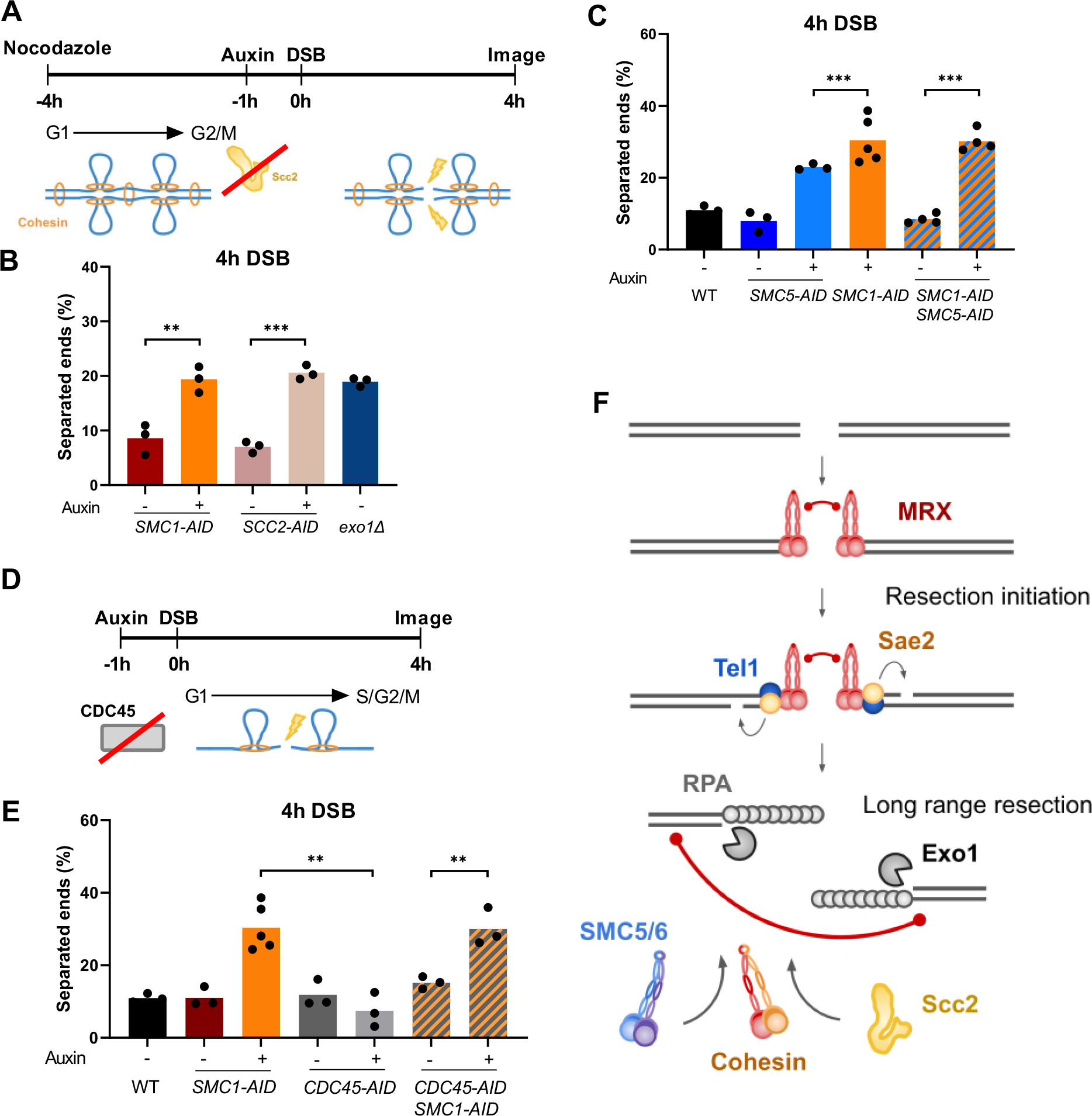
Cohesin DSB end-tethering requires de novo cohesin loading but not sister chromatid cohesion. **(A)** Schematic representation of assay to determine DSB end-tethering in absence of *de novo* cohesin loading. DSB was induced after cells were blocked in G2/M with nocodazole for 3 hours, and incubated with auxin or ethanol for a further 1 hour. (B) Percentage of G2/M blocked cells with separated ends in the indicated strains after 4 hours DSB induction. (C) Percentage of cells with separated ends in the indicated strains after 4 hours DSB induction. (D) Schematic representation of assay to determine DSB end-tethering in absence of replication. Cultures were incubated with auxin or ethanol for 1 hour. In the absence of Cdc45, cells advance through the cell cycle upon DSB induction, and load cohesin onto chromosomes without undergoing replication. (E) Percentage of cells with separated ends in the indicated strains after 4 hours DSB induction. (F) Schematic representation of DSB end-tethering pathways. Black stars indicate statistical differences (^*^ = p<0,05; ^**^ = p<0,01; ^***^ = p<0,005; ^****^ = p<0,001).

Previous studies have shown the importance of Smc5/6 in enriching cohesin at DSBs (*14*). To further explore this, we depleted Smc5 in our DSB end-tethering assay (fig. S2C). At 4 hours post-DSB induction, Smc5 depletion resulted in a significant increase in DSB end separation (Fig. 2C). Simultaneous depletion of Smc5 and Smc1 did not increase end separation beyond that observed upon Smc1 depletion alone (Fig. 2C), indicating that cohesin and Smc5/6 function in the same DSB end-tethering pathway. In conclusion, at 4 hours post-DSB, *de novo* cohesin loading at DNA DSB sites, mediated by Scc2/4 and Smc5/6, is necessary for DSB end-tethering.

### Cohesin DSB end-tethering does not require sister chromatid cohesion

Despite efficient cleavage of both sister chromatids in our assay (fig. S1A), which makes tethering of a cleaved chromatid by its sister unlikely, we aimed to confirm that DSB end-tethering was independent of sister chromatid cohesion. In absence of Cdc45, G1 cells progress to G2/M phase and load cohesin on chromosomes without firing replication origins and synthesizing sister chromatids (*31*), enabling us to assess the role of cohesin in DSB end-tethering in the absence of sister chromatid cohesion (Fig. 2D, and fig. S2D and S3A). Depletion of Cdc45 did not disrupt DSB end-tethering at 4 hours post-DSB induction (Fig. 2E), indicating that the presence of a sister chromatid is not essential for DSB end-tethering. Additional depletion of Smc1 resulted in increased DSB end separation, reaching levels similar to those observed in cells depleted of Smc1 alone. This indicates that cohesin can tether DSB ends even in the absence of DNA replication and a sister chromatid.

Together, these findings unveil a series of events that ultimately result in cohesin-dependent DSB end-tethering (Fig. 2F). Initially, the MRX complex binds and tethers DSB ends. Later, an Exo1-dependent pathway comes into play with the recruitment and *de novo* loading of cohesin at DSBs, facilitated by Scc2/4 and Smc5/6, actively participating in the tethering of DSB ends within individual chromatids.

### Cohesin orchestrates compaction of DSB flanking chromatin

Cohesin has been shown to form DNA loops and we hypothesized this activity could contribute to DSB end-tethering. To gain insights into the behavior of cohesin in the chromatin surrounding DSB, we modified our DSB end-tethering system to investigate chromatin compaction in a 48 kb region flanked by LacO-LacI-mCherry and TetO-TetR-GFP arrays, located 7 kb upstream of the DSB site (Fig. 3A). We measured the distance between these two signals in the presence or absence of DSB, to evaluate DSB-induced chromatin compaction (Fig. 3B). As the occurrence of a DSB triggers the DNA damage checkpoint and a G2/M cell cycle arrest, we treated all cells with nocodazole to ensure a fair comparison between DSB and no-DSB conditions.

**Fig. 3.**
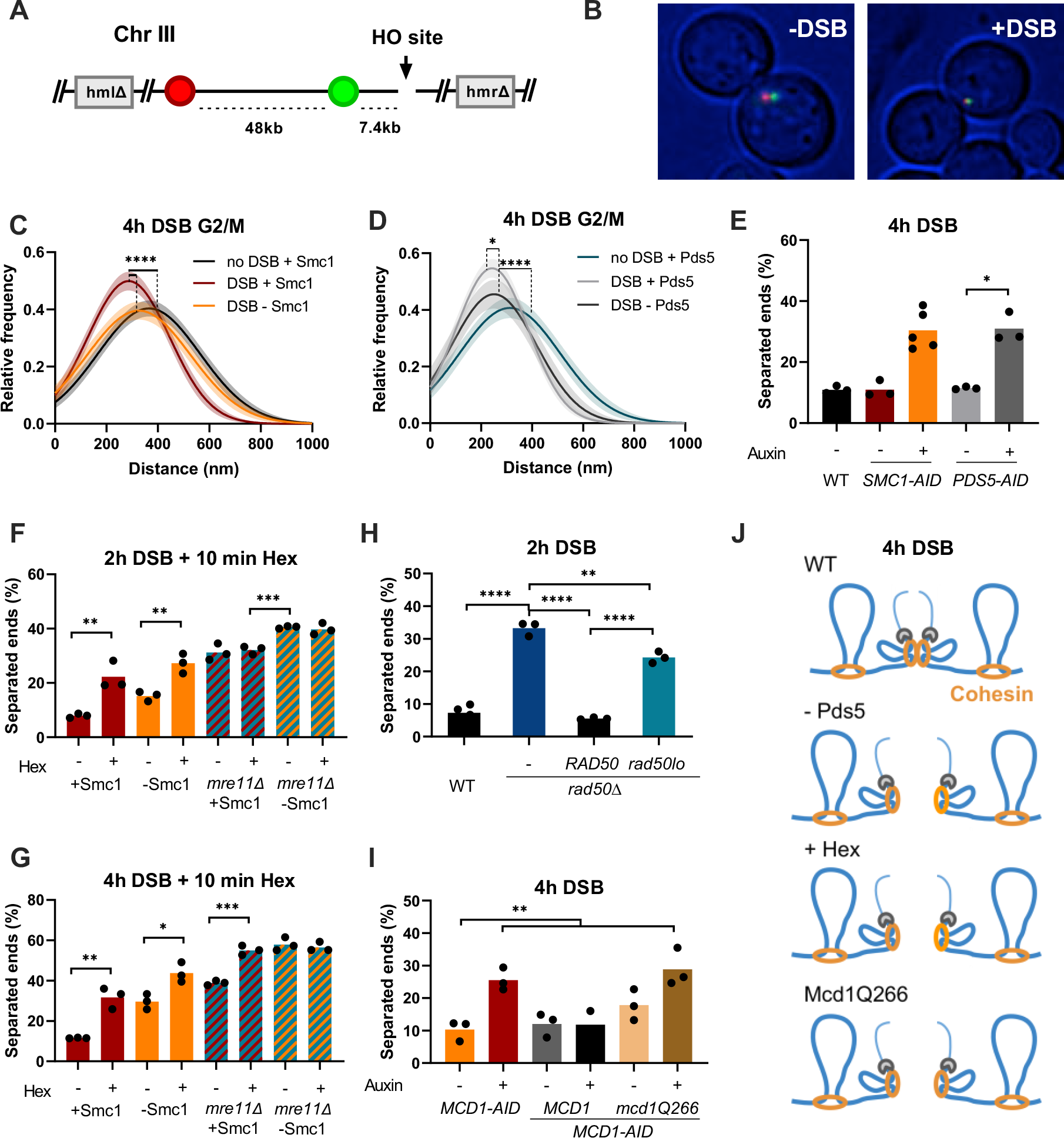
Cohesin compacts DSB flanking chromatin and DSB-ends are tethered by MRX- or cohesin-oligomerization. **(A)** LacO/LacI-mCherry tags and a TetO/TetR-GFP tag inserted at 7 and 55 kb from the HO DSB site at the MAT locus respectively. (B) Representative images in presence and absence of DSB. (C) Relative frequency of distances between the two tags in nocodazole arrested *SMC1-AID* tagged cells treated with ethanol (+Smc1) or auxin (-Smc1) after 4 hours and no DSB. (D) Relative frequency of distances measured between the two tags in nocodazole arrested *PDS5-AID* tagged cells treated with ethanol (+Pds5) or auxin (-Pds5) in after 4 hours or no DSB induction. (E)Percentage of cells with separated ends in the indicated strains after 4 hours DSB induction. (F-G) Percentage of cells with separated ends in the indicated strains treated with auxin or ethanol, and for 10 minutes with digitonin (-) or digitonin and 1,6-hexanediol (+), after 2 hours (F)or 4 hours (G) DSB induction. (H) Percentage of cells with separated ends in WT, rad50Δ, and rad50Δ cells complemented with *RAD50* or *rad50-lo*, after 2 hours DSB induction. (I) Percentage of cells with separated ends in MCD1-AID, and MCD1-AID strains complemented with *MCD1* or *mcd1-Q266*, in absence (-) or presence (+) of auxin, after 4 hours DSB induction. (J) Schematic representation of how loss of oligomerization disrupts DSB end-tethering. Black stars indicate statistical differences (^*^= p<0,05; ^**^= p<0,01; ^***^= p<0,005; ^****^= p<0,001). Shaded area indicates the 95% confidence interval of the fitting of 3 experiments per data set.

We first examined the impact of cohesin on chromatin folding in G2/M-arrested cells with no DSB. We observed a significant increase in the distribution of the distances upon cohesin depletion (fig. S4, A and B), showing that our assay enables detection of the previously reported cohesin-dependent compaction of chromatin (*32, 33*). Following 4 hours of DSB induction, we detected a significant reduction in distances between the two signals compared to the no-DSB condition, indicative of a compaction of the DSB-flanking chromatin (Fig. 3C; black versus red). This DSB-induced compaction was abolished upon depletion of Smc1, demonstrating that cohesin is responsible for the compaction of DSB flanking sequences (Fig. 3C; orange, and fig. S4C).

### Pds5 is required for DSB end-tethering but not DSB-induced genome compaction

If loop formation were at the basis of DSB end-tethering, the latter should be challenged by modulating loop expansion and turn over. To explore this, we tested the role of Pds5, a key factor responsible for cohesin loop regulation. Pds5 depletion weakens loop boundaries, reduces defined chromosome contacts/loops, and generate much longer loops in regions such as those near centromeres (*34, 35*). We found that DSB induced chromatin compaction still occurs in absence of Pds5 (Fig. 3D, and fig. S4, D-F). In contrast, Pds5 depletion increased end separation at 4 hours post-DSB, mimicking the effects of cohesin depletion (Fig. 3E). These results imply that either the loops formed in absence of Pds5 were not sufficient to support the function of cohesin in DSB end-tethering, or that cohesin tethers DSB ends independently of loop formation, through another mechanism requiring Pds5. A recent study has revealed an essential role of Pds5 in the oligomerization of multiple cohesin complexes (*36*), opening the door for a role of Pds5-dependent cohesin oligomerization in DSB end-tethering.

### Cohesin and MRX tethering rely on weak hydrophobic interactions

To investigate if protein-protein interactions and the oligomerization of cohesin complexes participates in DSB end-tethering, we used the aliphatic alcohol 1,6-hexanediol. Hexanediol has been instrumental in studying the liquid phase separation and oligomerization properties of various proteins, including cohesin and proteins involved in the DNA damage response (such as Rad52 and RPA; fig. S5, A and D-E; (*37, 38*)).

The treatment of cells with hexanediol 10 minutes prior to imaging at 2 hours post-DSB, when tethering mostly relies on MRX, increased end separation independently of Smc1 presence (Fig. 3F). Moreover, end separation was not increased by hexanediol in the absence of Mre11 alone. These results suggest a role for weak hydrophobic interactions in MRX-dependent tethering. In addition, hexanediol had no effect in cells depleted for both Mre11 and Smc1, suggesting that no other weak hydrophobic interactions intervene in DSB end-tethering at this early stage. Strikingly, hexanediol-treated *mre11Δ* cells do not exhibit the separation levels observed in Smc1 depleted *mre11Δ* cells (Fig. 3F, and fig. S1C), with or without hexanediol treatment (Fig. 3F). This finding aligns with a recent *in vivo* study in *S. cerevisiae* that demonstrated the resistance of a subset of topologically important cohesins to hexanediol treatment (*37*). As hexanediol is known to disrupt protein-protein interactions, this further supports our early finding that an Exo1-independent population of cohesin can tether DSB ends that are formed within a cohesin loop (fig. S1, C and D, and fig. S5F).

At 4 hours post-DSB, hexanediol-treated control cells also exhibited untethering (Figure 3G). In line with our observation at 2 hours that hexanediol disrupts MRX-dependent tethering, hexanediol and Smc1-depletion have additive effects at 4 hours. Hexanediol also increases end separation in absence of Mre11 suggesting that it also disrupts cohesin-dependent DSB end-tethering. Strikingly, in contrast to the 2-hour time point, hexanediol increased end separation in *mre11Δ* cells to levels comparable to cells depleted for both Smc1 and Mre11 (Fig. 3G). These results indicate that protein-protein interactions play a key role in DSB end-tethering by both MRX and cohesin (Fig. 3J).

MRX has been shown to form oligomers *in vitro* and disruption of these oligomers by a mutation of the hydrophobic interaction patch within the Rad50 head domain (*rad50lo* mutant, (*39*)) led to the disappearance of DSB-dependent Mre11 foci *in vivo*. Since hexanediol also disrupts Mre11-GFP foci formation in our strain background (fig. S5, B and C), we introduced this mutation in our tethering system. Strikingly, complementation of *rad50Δ* cells with *rad50lo*, was unable to restore end-separation to WT levels at 2 hours post-DSB, unlike wild-type *RAD50* (Fig. 3H). Therefore, disrupting Rad50 head oligomerization impairs DSB end-tethering.

The cohesin subunit Mcd1 has been identified as a mediator of cohesin oligomerization, and a 5 amino-acid insertion at Q266 in its regulation of cohesion and condensation (ROCC) domain has been shown to abolish cohesin oligomerization potential *in vivo* (*36, 40*). To test the role of cohesin oligomerization in DSB end-tethering, we complemented *MCD1-AID* cells with the *mcd1-Q266* mutant in both our compaction and end-tethering strains (fig. S6, A and B). Critically, *mcd1Q266* mutants exhibited strong DSB-dependent genome compaction (fig. S6, C-H), indicating cohesin is recruited to DSB sites and able to form chromatin loops. However, unlike complementation with *MCD1, mcd1Q266* failed to restore DSB end-tethering to WT-like levels (Fig. 3I), confirming the importance of cohesin oligomerization in DSB end-tethering. Taken together, these results indicate that both MRX and cohesin employ an oligomerization-dependent mechanism to tether DSB ends (Fig. 3J).

### Cohesin is required for efficient DNA DSB repair by homology directed mechanisms

Having identified a role for cohesin in tethering DSB ends, we questioned its significance for repair. We took advantage of our tethering system, which contains direct homologous repeats flanking the inserted LacO and TetO arrays (Fig. 4A). Following DSB induction and resection initiation, progressive formation of ssDNA away from the DSB causes loss of the dsDNA substrate which is necessary for the binding of the LacI-mCherry and TetR-GFP fusion proteins, and gradually leads to the disappearance of the fluorescent signals. Resection also unmasks the direct homologous repeats, which can anneal and be used to resynthesize the broken DNA strand. This restores chromosome continuity but results in loss of the genetic material that previously separated the homologous repeats used for repair. Following resynthesis, either the red or the green signal reappears, depending on the repeats used for repair (Fig. 4, A and B). After completion of the repair process, cells are released from the DNA damage checkpoint and proceed through cell division (Fig. 4B).

**Fig. 4.**
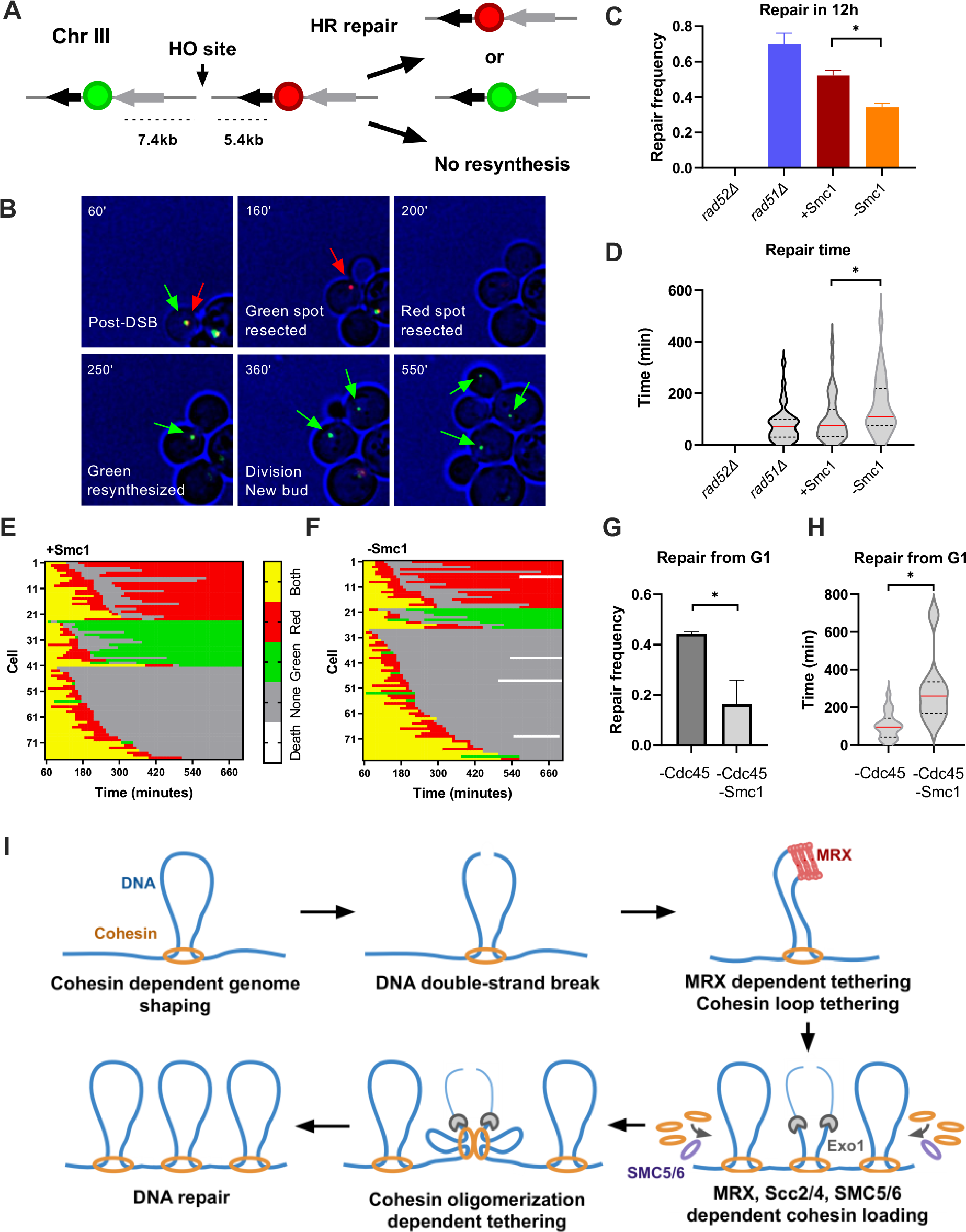
Cohesin is required for efficient DNA DSB repair by homology directed mechanisms. **(A)**Schematic representation of repair events after resection and disappearance of the spots followed by resynthesis of one spot. Black and grey triangles show direct repeats used for homologous recombination. (B) Sequence of images showing disappearance of both spots upon resection and reappearance of a green spot that is propagated to daughter cells at each division. Time post DSB is indicated on each frame. (C) Relative frequency of repair events corresponding to the resynthesis of a spot in *rad52Δ, rad51Δ* and *SMC1-AID* strains treated with ethanol (+Smc1) or auxin (-Smc1). (D) Time taken for a spot to reappear, in *rad52Δ, rad51Δ* and *SMC1-AID* strains treated with ethanol (+Smc1) or Auxin (-Smc1). (E-F) Spot characteristics of + Smc1 (C), and - Smc1 (D) individual cells imaged every 10 minutes during 12 hours after DNA DSB induction. Lines represent individual cell lineages, and each segment a time point. Colors indicate presence of both spots (yellow), a red spot only (red), a green spot only (green), or no spots (grey). (G) Relative frequency of repair events corresponding to the resynthesis of a spot in the indicated strains treated with auxin. Cells in G1 phase upon induction were imaged. (H) Time for a spot to reappear in the indicated strains treated with auxin. Cells in G1 phase upon induction were imaged. (I) Schematic representation of MRX and cohesin tethering DSB ends. MRX requires oligomerization through Rad50 head domains, and interaction between Rad50 coiled coils. Exo1 drives long-range DNA resection, leads to the recruitment of cohesin that mediates DSB end-tethering by oligomerization. Black stars indicate statistical differences (^*^= p<0,05; ^**^= p<0,01; ^***^= p<0,005; ^****^= p<0,001).

To assess repair events, we employed a microfluidics system to follow individual cells and image each fluorescent signal over a 12-hour period after DSB induction. To validate our assay, we imaged cells lacking *RAD52*, which is essential for all homology directed repair (HDR) events. In the absence of Rad52, no instances of spot reappearance were observed (Fig. 4C, and fig. S7A). Conversely, the loss of Rad51, which impedes gene conversion and promotes single-strand annealing (SSA), led to an increase in repair events compared to WT-like condition (SMC1-AID without auxin, Fig. 4, C and E, and fig. S7B), as previously reported (*41, 42*). This result suggests that inhibiting gene conversion, and favoring repair by SSA, leads to more detectable repair events in this assay, with unaltered repair kinetics compared to the WT-like condition (Fig. 4D). In contrast, upon Smc1 depletion, we observed a significant reduction in the frequency of repair events associated with a noticeable delay in repair kinetics (Fig. 4, C, D, E and F). This decrease in repair frequency was not caused by a resection defect (fig. S7C). To separate the dependence of repair events on sister chromatid cohesion from DSB end-tethering, we employed Cdc45 depletion. Strikingly, despite repair events still taking place upon Cdc45 depletion, simultaneous depletion with Smc1 resulted in a severe decrease in both the frequency of repair events and their kinetics compared to cells depleted of Cdc45 alone (Fig. 4, G, H and fig. S7, D-F). This indicates that the specific function of cohesin in DSB end-tethering is essential for efficient repair between DSB ends.

## Discussion

Cohesin enrichment at DSBs has long been known (*11*–*13*) with early studies also highlighting the importance of cohesin for survival after DNA damage inducing radiation (*11, 13, 43*). Recent observations have suggested that loop extrusion at DNA DSBs helps establish DNA damage response related chromatin modifications (*20*), which ultimately organize DSBs into microdomains (*44*). Moreover, sister chromatid cohesion, which is increased in response to DSB (*18, 19, 21*–*23*), prevents promiscuous repair events with far loci (*24, 27*).

In addition to these functions, we reveal a new cohesin role in tethering DSB ends. Cohesin’s first contribution, early after DSB formation is independent of MRX and Exo1 and likely relies on cohesin-dependent genome looping, as predicted by recent theoretical work (*2*). Later, cohesin-dependent DSB end-tethering requires *de novo* cohesin loading, acts in cooperation with Exo1 and Smc5/6, is independent of sister chromatid cohesion and loop formation, and relies on cohesin oligomerization. Importantly, our data provide a biological function to the recently described cohesin oligomerization mechanism (*36, 37*) that is independent of cohesin’s canonical roles in sister chromatid cohesion and loop extrusion.

Strikingly, our results support the existence of two populations of DSB-bound cohesin with separable functions, namely chromatin compaction and DSB end-tethering, and different modes of action, namely loop formation and oligomerization. One population of cohesin acts in a Pds5- and oligomerization-independent manner and compacts DSB adjacent chromatin. This cohesin-dependent compaction may participate in DSB signaling though a loop extrusion-mediated spreading of histone H2AX phosphorylation, as previously suggested (*20*). A second population requires Pds5 and cohesin oligomerization, and tethers DSB ends. What distinguishes loop-forming cohesin from DSB end-tethering cohesin, beyond the capacity to form oligomers, is unknown. However the fact that the DSB end-tethering cohesin population acts independently of MRX, which has been implicated in cohesin enrichment at DSBs (*12, 15*), suggests a new mode of recruitment of these cohesin to DSB ends.

Our data supports a role for Scc2, Smc5/6 and Exo1 mediated ssDNA formation in recruiting or stabilizing DSB end-tethering cohesin. Whereas Scc2 and Smc5/6 were previously implicated in the recruitment of cohesin to DSB, the formation of ssDNA by Exo1 is specifically required for cohesin-dependent DSB end-tethering. Since dsDNA bound cohesin can capture ssDNA (*45*), formation of ss-DNA may directly intervene in cohesin recruitment. Bridging dsDNA with ssDNA could also be sufficient to account for DSB end tethering. Otherwise, cohesin recruitment could be mediated by Smc5/6, which interacts with ssDNA through its hinge domain (*46, 47*), and stably associates with ss-dsDNA junctions (*47, 48*). Smc5/6, that bears both ubiquitin and SUMO ligase activity, could then locally modify a pool of cohesin, promoting cohesin oligomerization and DSB end-tethering.

Our results, which reveal cohesin’s role in DSB end-tethering, contrast with a previous report suggesting that cohesin is dispensable for contacts between both sides of a DSB as captured by a Hi-C approach (*24*). One plausible explanation for this discrepancy is rooted in the technologies used to make such observations. Single-cell live-microscopy is more sensitive at this scale considering we detect DSB-induced compaction beyond G2/M and cohesin-dependent loss of end tethering, both appearing below the detection threshold of the population-wide Hi-C approach (*24*).

We also show that Rad50 (MRX^MRN^) head oligomerization is required for MRX dependent DSB end-tethering. MRX oligomerization via both the Rad50 heads and coiled coils has been described in both yeast and humans (*39, 49*). Our data demonstrates that Rad50 head oligomerization observed *in vitro*, is significant for MRX end-tethering *in vivo*. Alongside the necessity of the Rad50 Zn-hook for DSB end-tethering demonstrated *in vivo* (*3*), our data supports the Velcro model recently proposed based on structures frequently observed by electron microscopy for MRX-driven DSB end-tethering (*49*).

Together, our results suggest that oligomerization of SMC complexes is a conserved and functionally relevant mechanism for maintaining genome integrity in response to DNA damage. Interestingly, hexanediol treatment disrupted MRX foci in response to DSB, suggesting MRX at DSBs may form condensates. Although, cohesin does not form detectable foci in response to DSB in yeast, it has been shown to form phase separation condensates *in vitro* (*37*). These observations question the relevance of phase separation in DSB end-tethering, which should be investigated using single molecule microscopy in the future.

Given the prevalence of chromosome translocations in cancer, and the role of DSB induction in cohesin sensitive developmental processes such as V(D)J recombination (*50*) our study gives further insights into how SMC complex dysregulation may lead to disease in the human population.

## Acknowledgements

We thank F. Ulhman, S. Marcand, A. Quinet, A. Campalans, P. Radicella and P. Bertrand for critical reading of this manuscript and members of the Dubrana and Marcand laboratories for stimulating discussions. We thank Douglas Koshland and Matthias Peter for sharing plasmids and strains.

This research was supported by the European Research Council under the European Community’s Seventh Framework Program (FP7/2007 2013/European Research Council Grant Agreement 281287), Fondation ARC pour la Recherche sur le Cancer (PJA-), CEA Radiation biology and Impulsion programs, EDF. JP was supported by a fellowship from the CEA.

## Authors Contributions

JP, MT, CD, CB and RC performed experiments. KD designed and supervised the entire project with the help of MT and JP. KD and JP analyzed the data, assembled the figures and wrote the manuscript with critical input of the other authors.

## Declaration of interests

The authors declare no competing interests

## List of supplementary material

Material and Methods

Fig. S1-S7

Tables S1-S3

References 28-29, 51-54

Movie S1-S4

## Notes

### Competing Interest Statement

The authors have declared no competing interest.

