## supplementary materials for "Cohesin complex oligomerization maintains end-tethering at DNA double-strand breaks"

**The PDF file includes:**

Materials and Methods
Figs. S1 to S7
Tables S1 to S3
References 28-29, 51-54

**Other Supplementary Materials for this manuscript include the following:**

Movies S1 to S4

### Materials and Methods

#### Strains and plasmids

Yeast strains used in this study are derivative of JKM179, JKM139 (*51*) or yKD809 (*28*), and were generated by PCR gene targeting, plasmid transformation or cross (Table S1-S2).

#### Media and growth conditions

Yeast strains were grown at 30°C in glucose rich yeast extract-peptone-dextrose (YPD) medium, with appropriate antibiotic, or in synthetic medium (SD) lacking the appropriate amino acid. YPLGg medium containing 2% lactate, 3% glycerol and 0.05% glucose was used for DNA DSB induction, by addition of galactose (f.c. 2%), to ON cultures of OD600 0.4 - 0.8 as in (*52*). Conditional protein knockdown was achieved in AID tagged strains by addition of IAA in EtOH to a f.c. of 2Mm (*29*) 1,6-hexanediol treatment (f.c. 10%) was performed for 10 minutes, with 10µg/ml digitonin.

#### Microscopy

Live-cell images were acquired using a wide-field inverted micro-scope (Leica DMI-6000B) equipped with Adaptive Focus Control to eliminate Z drift, a 100×/1.4 NA immersion objective with a PriorNanoScanZ Nanopositioning Piezo Z Stage System, a CMOS camera(ORCA-Flash4.0; Hamamatsu) and a solid-state light source (Spec-traX, Lumencore). The system is piloted by MetaMorph software(Molecular Device). Images were acquired at indicated time points after DSB induction. 19 focal steps of 0.20µm were acquired sequentially for GFP and mCherry with an exposure time of 50ms using solid-state 475- and 575-nm diodes and appropriate filters (GFP-mCherry filter; excitation: double BP, 450–490/550–590 nm and dichroic double BP 500–550/600–665 nm; Chroma Technology Corp.). Images were processed using ImageJ software (National Institutes of Health). 3D images were converted to 2D projections, from which XY coordinates of the most intense pixels were extracted. Distance analysis between proximal fluorescent signals in mCherry and GFP channels was performed using an Rstudio script. All images shown are z projections of z-stack images.

#### Quantifications and statistical analysis

Quantifications and statistical analysis were done using PRISM (GraphPad). For the end-tethering assay, at least 3 independent experiments analysing more than 100 cells were performed for each genotype and statistical significance was determined by a two-tailed Student's t test. ns=not significant p≥0,005, * = p<0,05; ** = p<0,01; *** = p<0,005; **** = p<0,001. For the compaction measurements, distance data of at least 100 cells was sorted into 200 nm bins, and the bins of 3 independent experiments were fitted with a gaussian curve using Prism software, with shaded areas representing an interval of confidence of 95%. Statistical significance was determined by a Kolmogorov-Smirnov test. ns=not significant p≥0,005, * = p<0,05; ** = p<0,01; *** = p<0,005; **** = p<0,001.

**Microscopy in microfluidic plates**

CellASIC ONIX microfluidic plates (Y04C-02; MilliporeSigma) were used for long duration movies. HO was induced in YPLGg cultures of OD600nm 0.5 by addition of galactose to a f.c. of 2%, and incubation at 30°C for 30 minutes. After break induction, cultures were loaded into the microfluidic plate. The remaining culture was centrifuged at 3000rpm for 3 minutes, and the conditioned media was loaded into the microfluidic plate for flow over the cells for the duration of the experiment. After loading the plate, cell positions were defined, and images were acquired every 10 minutes for up to 24 hours. 19 focal steps of 0.20µm were acquired sequentially for GFP and mCherry with an exposure time of 30ms using solid-state 475- and 575-nm diodes and appropriate filters (GFP-mCherry filter; excitation: double BP, 450–490/550–590 nm and dichroic double BP 500–550/600–665 nm; Chroma Technology Corp.). A single bright-field image on one focal plane was acquired at each time point with an exposure of 10ms. For Cdc45 depleted strains, cells were loaded into the microfluidic plate immediately following galactose addition, and cells that were in G1 prior DSB induction were imaged.

**Monitoring DSB efficiency**

Cells were grown in 2ml of YPD ON. Cultures were then diluted in YPLGg, and grown to an OD600nm of 0.5-0.8, and incubated with 2mM IAA or EtOH for 1 hour. HO expression was induced by addition of galactose to a final concentration of 2%. At 0, 1, 2, 4 and 6 hours post DSB induction, approximately 4 x $10^7$ cells were collected by 3000rpm centrifugation for 5 min. DNA was extracted from cell pellets by Winston preparation. Samples were analyzed by qPCR with primers 1kb upstream of the HO site to analyze resection (200nM), flanking the HO site to determine DSB efficiency (450nM) or targeting the OGG1 reference gene (200nM). See Table 3 for primer sequences. Reactions were performed as in (*53*). Each sample and no template controls were run in triplicate, and reaction specificity determined by melt curve analysis. Relative quantitation of resection and DSB efficiency reactions was achieved using the comparative Ct method (*54*).

**Western blot**

Auxin induced protein degradation of AID containing strains was confirmed by Western blot analysis (Brocas et al., 2019). Cells were grown in 2ml of YPD ON. Cultures were then diluted in YPLGg, and grown to an OD600nm of 0.5-0.8, and incubated with 2mM IAA or EtOH for 1, 2, and 4 hours (hrs). Approximately 4 OD600nm of cells were collected by centrifugation at 3000rpm for 5 min. Cells were washed in dH2O, and collected by centrifugation at 3000rpm for 5 min. Supernatant was removed, and cell pellets frozen at -80°C. Whole cell extracts were prepared from cell pellets using a standard Trichloroacetic acid (TCA) extraction protocol and suspended in Laemmli buffer. Protein concentrations were determined by Bradford assay, and samples prepared for SDS PAGE by 5 min incubation at 90°C. 20µg of sample was migrated at 100v for 1 hr on 10% SDS polyacrylamide gels in standard running buffer. Nitrocellulose membrane transfer was performed using the iBlot transfer apparatus as per manufacturers guidelines (Thermo Fisher).

Membranes were washed with TBS-T, revealed by ponceau staining, and blocked with 5% milk TBS-T for 1hr. Membranes were then incubated at room temperature with mouse primary anti-myc (1:1000), and anti-mouse secondary antibodies (1:1000) in 5% milk TBS-T for 1hr each. Membranes were developed by fluorescence using the Odyssey Clx (LI-COR).

**Flow cytometry**

0.5 OD600nm of cells were fixed in ethanol 70% and stored at -20°C. Cells were pelleted, washed, and then incubated in Sodium Citrate pH7.4 50mM with 0.25mg/ml RNAseA for 1 hour at 50°C. Proteinase K was then added, to a final concentration of 2mg/ml, and incubated for further 1 hour at 50°C. Cells were pelleted, and then stained in a pH7.4 50mM Sodium Citrate solution containing 1μM SYTOX Green Nucleic Acid Stain (Invitrogen – S7020). Cells were sonicated, and flow cytometry was performed on a Novocyte cytometer (ACEA bioscience.Inc). Data was analyzed using FlowJo software.

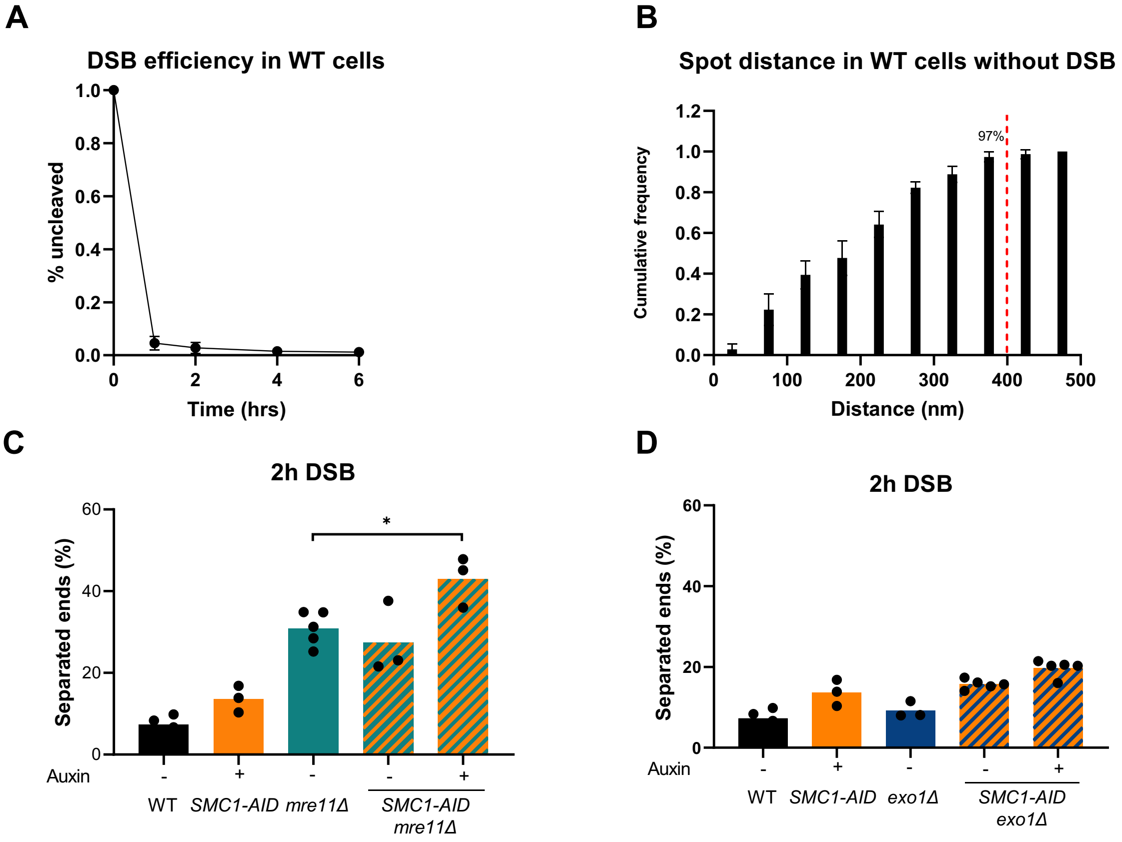


**Fig. S1. LacI-mCherry and TetR-GFP spot distance rarely exceed 400nm, and HO induced DSB is fast and efficient. Cohesin contributes to DSB end tethering at 2 hours post-DSB in absence of MRX.**

(A) qPCR detection of the HO cleavage site in WT cells at 0, 1, 2, 4 and 6 hours after DSB induction. (B) Cumulative distance between LacI-mCherry and TetR-GFP signals in exponential WT cells without DNA DSB induction. Red line indicates 400nm threshold, which 97% of distances are under. (C) Percentage of cells with separated ends in WT, SMC1-AID, mre11Δ, mre11Δ SMC1-AID strains after 2h DSB induction. (D) Percentage of cells with separated ends in WT, SMC1-AID, exo1Δ, exo1Δ SMC1-AID strains after 2h DSB induction.

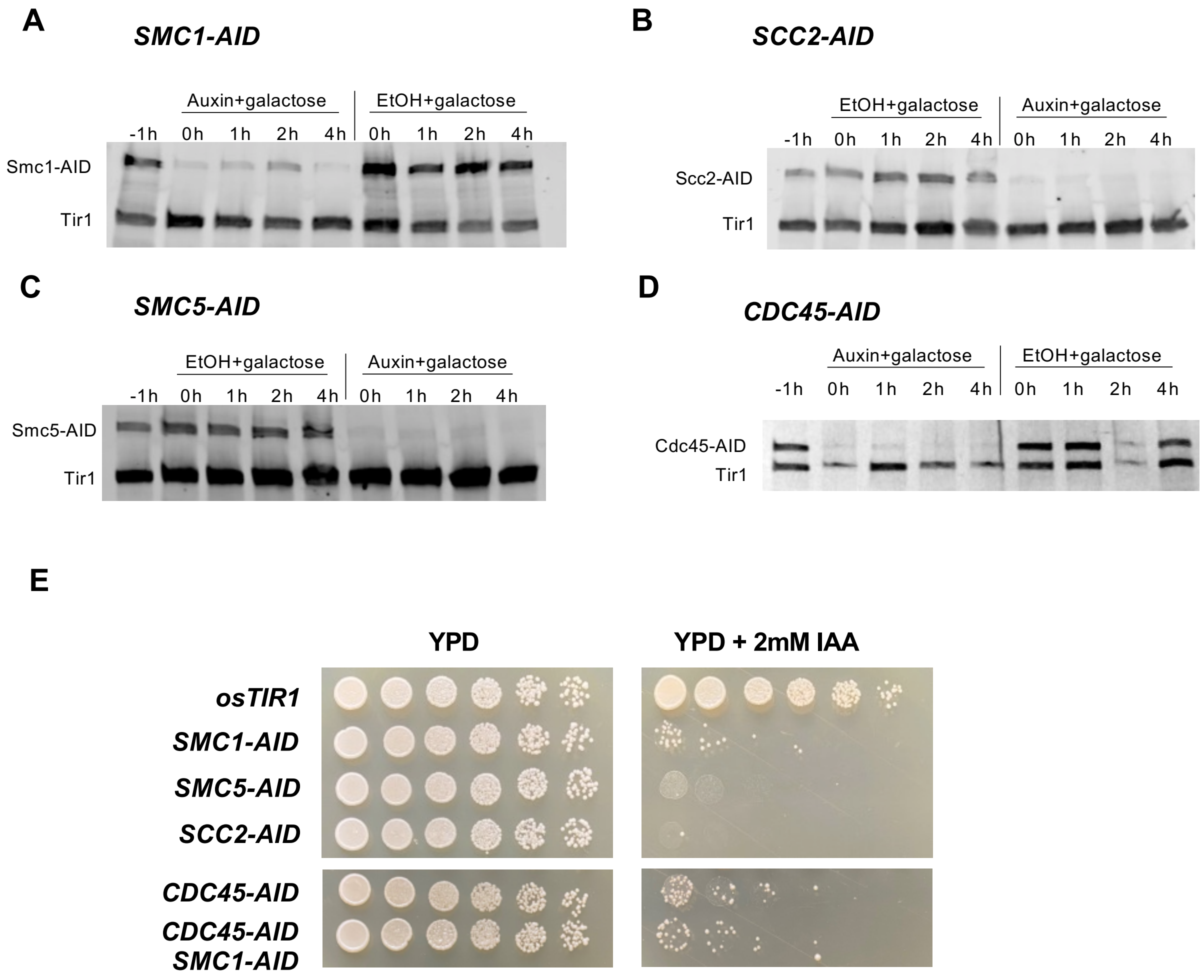


**Fig. S2. Auxin induced degradation of target proteins leads to efficient depletion.**

(A-D) anti-myc Western blots demonstrating protein levels of 9myc-AID tagged proteins treated with auxin or ethanol throughout microscopy DSB end tethering assays. t-1 (before IAA/EtOH addition), t0 (1 hour IAA/EtOH), t1 (2 hour IAA/EtOH + 1h galactose), t2 (3 hour IAA/EtOH +2h galactose) and t4 (5 hours IAA/EtOH + 4h galactose). (E) Drop assay of all tethering strains on YPD and YPD + auxin, incubated for 48 hours at 30°C.Type or paste caption here.

A

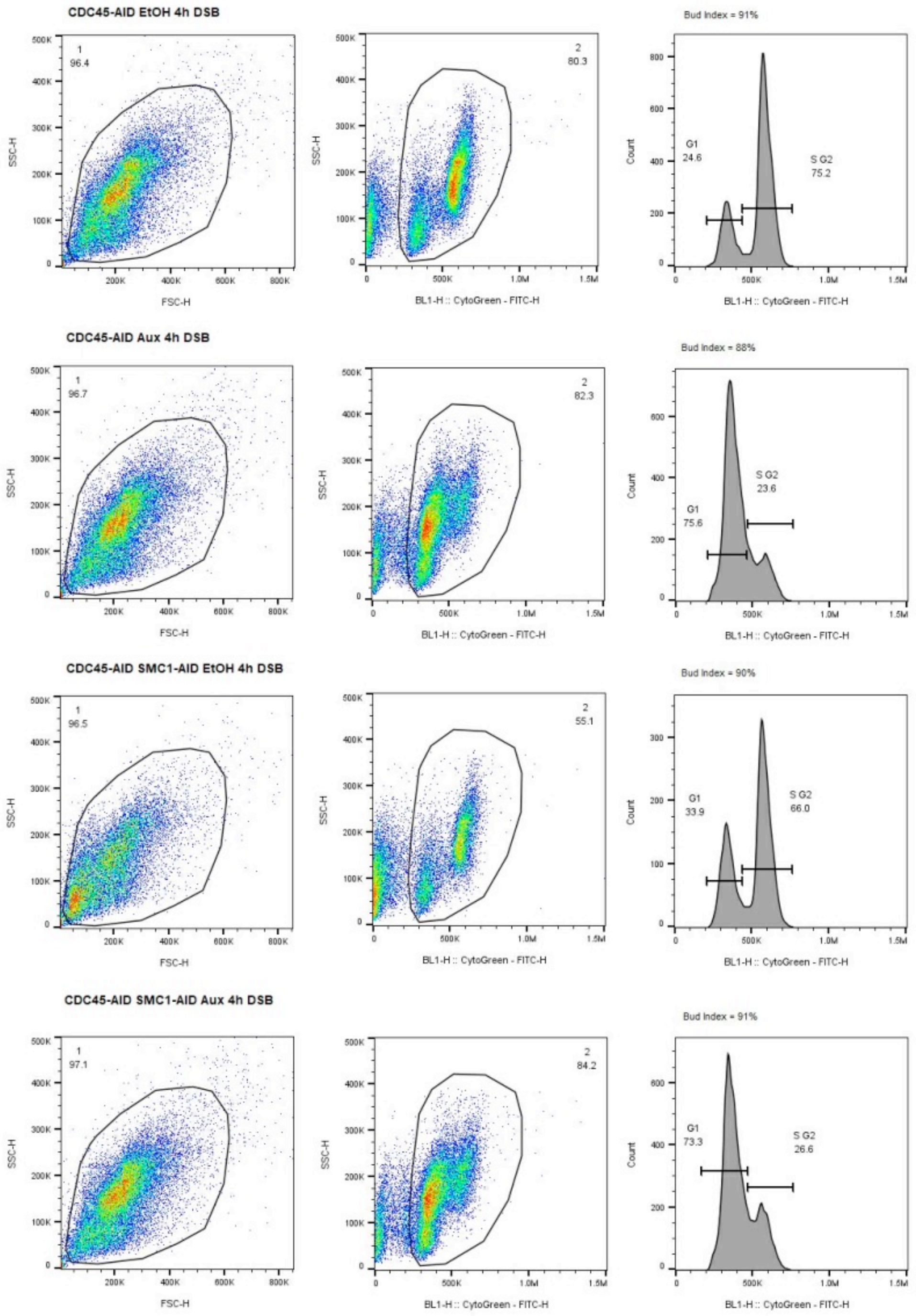


**Fig. S3. Cdc45 degradation prevents genome duplication whilst allowing cells to proceed to G2/M.**

(A) Gating and fluorescent intensity profiles, determind by flowcytometry, of CDC45-AID and CDC45-AID SMC1-AID strains treated with IAA or EtOH after 4h DSB induction. Percentage of cells with large buds is indicated above intensity profiles as bud index.

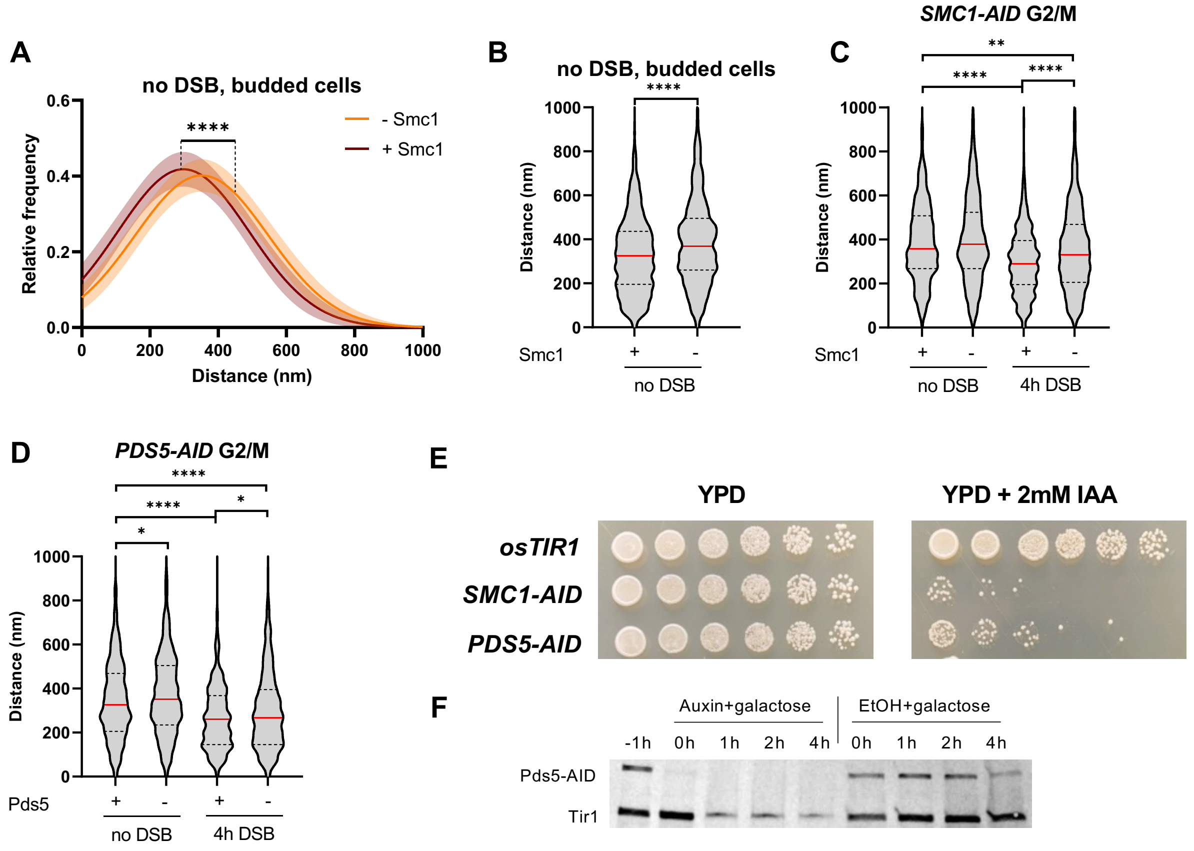


**Fig. S4. Smc1 depletion reveals cohesin dependent genome compaction in S-M phase cells. Pds5 is not required for DSB dependent genome compaction.**

(A) Relative frequency of distances measured between two tags separated by 45kb in a SMC1-AID tagged strain treated with ethanol (+SMC1) or auxin (-SMC1) in cycling cells in which a bud is present. (B) Distances between 45kb separated tags from three individual replicas for SMC1-AID tagged strain treated with ethanol (+Smc1) or auxin (-Smc1), represented as a violin plot. Red line at median, quartiles represented by dashed line. (C-D) Distances between 45kb separated tags from three individual replicas for SMC1-AID and PDS5-AID tagged strains treated with noodazole and following 4 hours DSB, treated with ethanol or auxin, represented as a violin plot. Red line at median, quartiles represented by dashed line. (E) Drop assay of compaction strains plated on YPD and YPD + auxin, incubated for 48 hours at 30°C. (F) anti-myc Western blot demonstrating protein levels of PDS5-AID strains treated with auxin or ethanol throughout microscopy DSB end tethering assay timecourse. t-1 (before IAA/EtOH addition), t0 (1 hour IAA/EtOH), t1 (2 hour IAA/EtOH + 1 hour galactose), t2 (3 hours IAA/EtOH +2 hours galactose) and t4 (5 hours IAA/EtOH + 4 hours galactose).

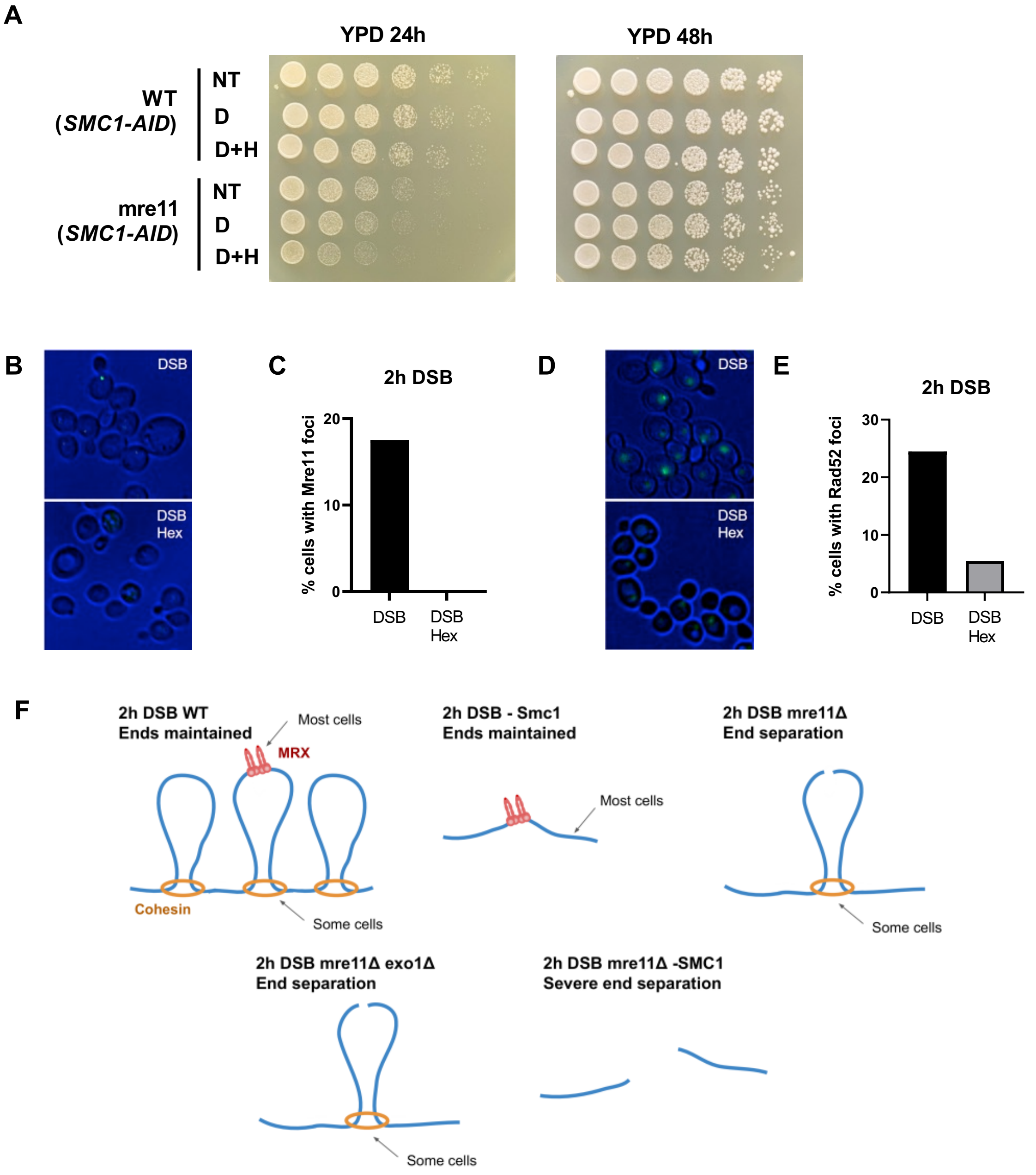


**Fig. S5. Cells recover following hexanediol treatment without growth defect, and Mre11-GFP foci are abolished by hexanediol treatment. A hexanediol resistant cohesin population reduces end separation in the absence of MRX at 2 hour DSB.**

(A) Drop assay of strains plated on YPD after no treatment (NT), 10 minutes digitonin (D), or 10 minutes digitonin + hexanediol (D+H) treatment, incubated for 24 and 48 hours at 30°C. (B) Representative images of Mre11-GFP foci at 2 hour DSB with no treatment (DSB), or 30 minutes digitonin + hexanediol (DSB Hex) treatment. (C) Quantification of cells with Mre11-GFP foci at 2 hour DSB with no treatment (DSB), or 30 minutes digitonin + hexanediol (DSB Hex) treatment. (D) Representative images of Rad52-YFP foci at 2 hour DSB with no treatment (DSB), or 30 minutes digitonin + hexanediol (DSB Hex) treatment. (E) Quantification of cells with Rad52-YFP foci at 2 hour DSB with no treatment (DSB), or 30 minutes digitonin + hexanediol (DSB Hex) treatment. (F) Schematic representation of cohesin dependent end tethering in a looping dependent manner. In WT cells, MRX tethers early DSB ends, and the DSB might be in a cohesin loop. Cohesin depletion following 2 hour DSB doesn't increase end separation due to MRX compensation. In the absence of MRX, DSB ends that occured within a cohesin loop rescues some loss of DSB end tethering. In contrast, loss of both MRX and cohesin leads to increased end separation as both the cohesin and MRX dependent mechanisms are lost.

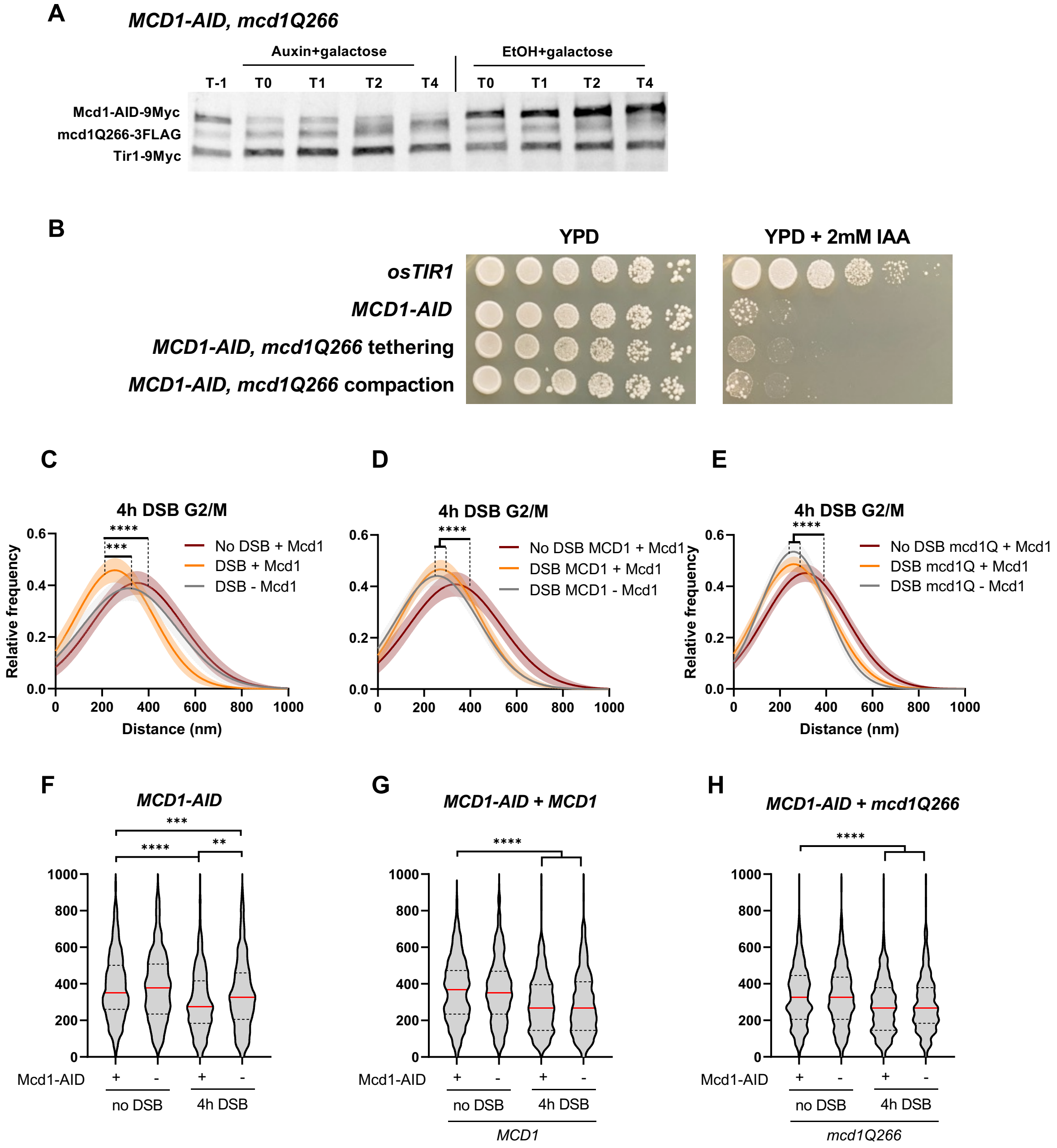


**Fig. S6. mcd1Q266 rescues DSB dependent genome compaction in the absence of Mcd1.**

(A) anti-myc/anti-flag Western blots demonstrating protein levels of AID-myc and mcd1Q266-FLAG tagged proteins treated with auxin or ethanol throughout microscopy DSB end tethering assays. t-1 (before IAA/EtOH addition), t0 (1 hour IAA/EtOH), t1 (2 hours IAA/EtOH + 1 hours galactose), t2 (3 hours IAA/EtOH +2 hours galactose) and t4 (5 hours IAA/EtOH + 4 hours galactose). (B) Drop assay of MCD1 strains on YPD and YPD + auxin, incubated for 72 hours at 23°C. (C-E) Relative frequency of distances measured between two tags separated by 45kb in MCD11-AID tagged strains complemented with nothing, MCD1, or mcd1Q266, treated with ethanol or auxin and nocodazole after 4h DSB. (F-H) Distances between 45kb separated tags from three individual replicas for *MCD1-AID* tagged strains complemented with nothing, *MCD1*, or *mcd1Q266*, treated with ethanol or auxin and nocodazole after 4 hour DSB, represented as a violin plot. Red line at median, quartiles represented by dashed line.

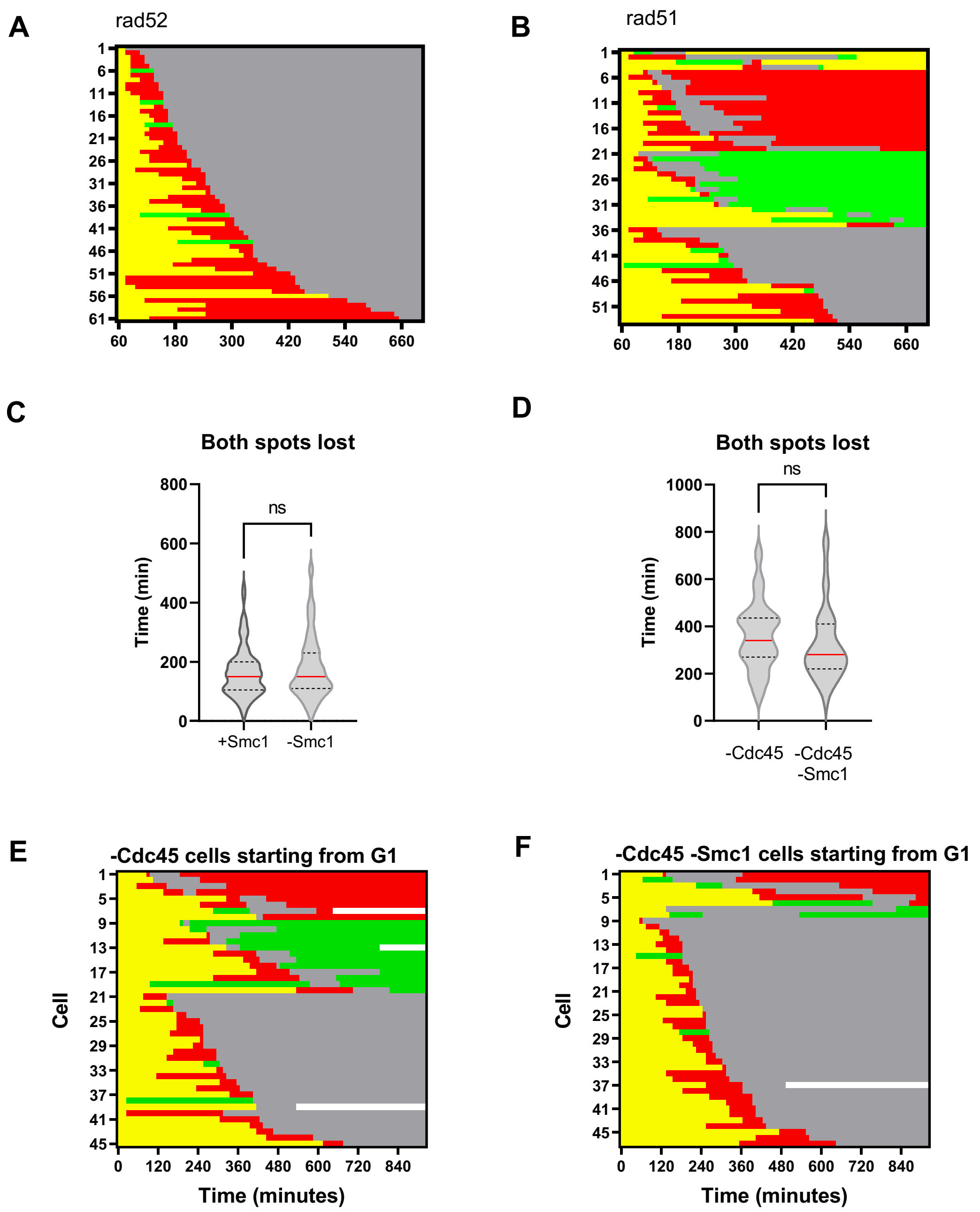


**Fig. S7. Cohesin depletion does not alter rate of resection following DSB.**

(A-B) Spot characteristics of individual cells in *rad52* and *rad51* cells during a 12 hour period after DNA DSB induction (C) Time taken for loss of both spots after DSB induction in microfluidic experiments for *SMC1-AID* strains. (D) Time taken for loss of both spots after DSB induction in microfluidic experiments for auxin exposed *CDC45-AID* and *CDC45-AID SMC1-AID* strains. (E-F) Spot characteristics of individual cells in *CDC45-AID*, *CDC45-AID SMC1-AID* cells during a 15.5 hour period after DNA DSB induction. DSB was induced in the microfluidic plate.

| Strain | Strain genotype | Strain type | Reference | Figure ID |
|---|---|---|---|---|
| JKM139 | *MATa hml::ADE1 hmr::ADE1 ade3::pGal-HO ade1 leu2-3,112 lys5 trp1::hisG ura3-52* | Galactose-inducible HO cleavage | Lee et al., 1998 | |
| JKM179 | *MATα hml::ADE1 hmr::ADE1 ade3::pGal-HO ade1 leu2-3,112 lys5 trp1::hisG ura3-52* | Galactose-inducible HO cleavage | Lee et al., 1998 | |
| yKD809 | JKM139, *ura3-52Δ::LacI-mCherry-URA3, leu2-3Δ::TetR-GFP-LEU2, TAF2-LacOpFx-TRP1, 4.4kb MATa-TetO-LEU2* | Tethering | Mojumdar et al. , 2019 | 1B-D, 1F-H, 2C, 2E, 3E, 3H, S1A, S1C-D |
| yKD1107 | yKD809*, exo1Δ::HPH* | Tethering | This study | 1C-D, 1H, 2B, S1D |
| yKD925 | yKD809, *mre11Δ::HPH* | Tethering | Mojumdar et al. , 2019 | 1C-D, 1G, S1C |
| yKD2365 | yKD809, *exo1Δ::HPH mre11Δ::Nat* | Tethering | This study | 1C-D |
| yKD1175 | yKD809, *ura3-52Δ:: OsTIR1-9myc-URA3-LacI-mCherry-KanMx* | Tethering | This study | S1B, S2E, S6B |
| yKD1177 | yKD1175, *SMC1-AID-9myc-HPH* | Tethering | This study | 1E-H, 2B-C, 2E, 3E-G, 4B-F, S1C-D, S2A, S2E, S5A, S7C |
| yKD2318 | yKD1175, *SMC1-AID-9myc-HPH mata-inc* | Tethering | This study | 1F |
| yKD1483 | yKD1175, *SMC1-AID-9myc-HPH mre11Δ::Nat* | Tethering | This study | 1G, 3F-G, S1C, S5A |
| yKD1486 | yKD1175, *SMC1-AID-9myc-HPH* exo1Δ::Nat | Tethering | This study | 1H, S1D |
| yKD1488 | yKD1175, *SCC2-AID-9myc-HPH* | Tethering | This study | 2B, S2B, S2E |
| yKD1178 | yKD1175, *SMC5-AID-9myc-HPH* | Tethering | This study | 2C, S2C, S2E |
| yKD2436 | yKD1175, *SMC1-AID-9myc-Nat SMC5-AID-9myc-HPH* | Tethering | This study | 2C |
| yKD2496 | *yKD1175, CDC45-AID-9myc-Nat* | Tethering | This study | 2E, 4G-H, S2D-E, S3A, S7D-E |
| yKD2497 | *yKD1175, CDC45-AID-9myc-Nat SMC1-AID-9myc-HPH* | Tethering | This study | 2E, 4G-H, S2E, S3A, S7D, S7F |

| | | | | |
|---|---|---|---|---|
| yKD2285 | *4.4kb MATa-TetO-LEU2 0.5kb-CWH43::lacOpFX-TRP1 ura3Δ::OsTIR1-URA3-LacI-mCherry-KanMx leu2-3Δ::TetR-GFP-LEU2* | Compaction | This study | S4E |
| yKD2289 | yKD2285, *SMC1-AID-9myc-HPH* | Compaction | This study | 3B-C, S4A-C, S4E |
| yKD2438 | yKD2285, *PDS5-AID-9myc-HPH* | Compaction | This study | 3D, S4D-F |
| yKD2439 | yKD1175, *PDS5-AID-9myc-HPH* | Tethering | This study | 3E |
| yKD2484 | yKD809, *ura3-52Δ::LacI-mCherry-KanMX rad50Δ::Nat* | Tethering | This study | 3H |
| yKD2549 | yKD2484, *ura3-52Δ::LacI-mCherry-RAD50-URA3* | Tethering | This study | 3H |
| yKD2550 | yKD2484, *ura3-52Δ::LacI-mCherry-rad50L116A/I119A/T127A/L128A-URA3* | Tethering | This study | 3H |
| yKD2485 | yKD1175, *ura3Δ::OsTIR1-Nat-LacI-mCherry-KanMx MCD1-AID-9myc-HPH* | Tethering | This study | 3I, S6B |
| yKD2491 | yKD2485, *mcd1-Q266-3FLAG-URA3* | Tethering | This study | 3I, S6B |
| yKD2492 | yKD2485, *MCD1-3FLAG-URA3* | Tethering | This study | 3I, S6B |
| yKD1172 | yKD809, *rad52Δ::KanMx* | Tethering | This study | 4C-D, S7A |
| yKD2366 | yKD809, *rad51Δ::Nat* | Tethering | This study | 4C-D, S7B |
| yKD2552 | yKD2285, *ura3Δ::OsTIR1-Nat-LacI-mCherry-KanMx* | Compaction | This study | S6B |
| yKD2553 | yKD2552, *MCD1-AID-9myc-HPH* | Compaction | This study | S6C, S6F |
| yKD2554 | yKD2553, *mcd1-Q266-3FLAG-URA3* | Compaction | This study | S6E, S6H |
| yKD2555 | yKD2553, *MCD1-3FLAG-URA3* | Compaction | This study | S6D, S6G |
| yKD1598 | JKM139, *NUP49::NUP49-mCherry-URA3 MRE11-yEGFP-HPH, sae2::NAT* | DDR foci | This study | S5B-C |
| yKD282 | JKM179, *RAD52-YFP* | DDR foci | This study | S5D-E |

**Table S1.**
*Saccharomyces cerivisae* strains used in this study

| Plasmid name | Plasmid number | Description | Source |
|---|---|---|---|
| OSTIR-9myc-URA3 | pKD243 | ADH1p-OsTIR1-9Myc | Nishimura et al., 2009 |
| AID-9myc-NAT | pKD244 | AID-9myc-NAT | Nishimura et al., 2009 |
| AID-9myc-HPH | pKD245 | AID-9myc-HPH | Nishimura et al., 2009 |
| mcd1-Q266-3FLAG-URA3 | pKD511 | pVG285 mcd1-Q266-3FLAG | Eng et al., 2014 |
| MCD1-3FLAG-URA3 | pKD517 | MCD1-3FLAG | This study |
| rad50lo-URA3 | pKD513 | pJT23 pSIVura-SpeI-prom-Rad50(L116A, I119A, T127A, L128A)-term-KpnI | Kissling et al., 2022 |
| RAD50-URA3 | pKD514 | pJT25, pSIVura-SpeI-prom-Rad50-term-KpnI | Kissling et al., 2022 |

**Table S2.**

Plasmids used in this study

| Primer | Name | Sequence | Use |
|---|---|---|---|
| 776 | F OGG1 | CAATGGTGTAGGCCCCAAAG | Reference gene |
| 777 | R OGG1 | ACGATGCCATCCATGTGAAGT | Reference gene |
| 1124 | F MATa | AGTTTCAGCTTTCCGCAACAG | Quantify DSB efficiency at MATa |
| 47 | R MATa | CGTCACCACGTACTTCAGCATAA | Quantify DSB efficiency at MATa |

**Table S3.**
qPCR primers used in this study

**Movie S1.**
yKD1177 (SMC1-AID) EthOH, cell repairs, resynthesises the green spot and divides

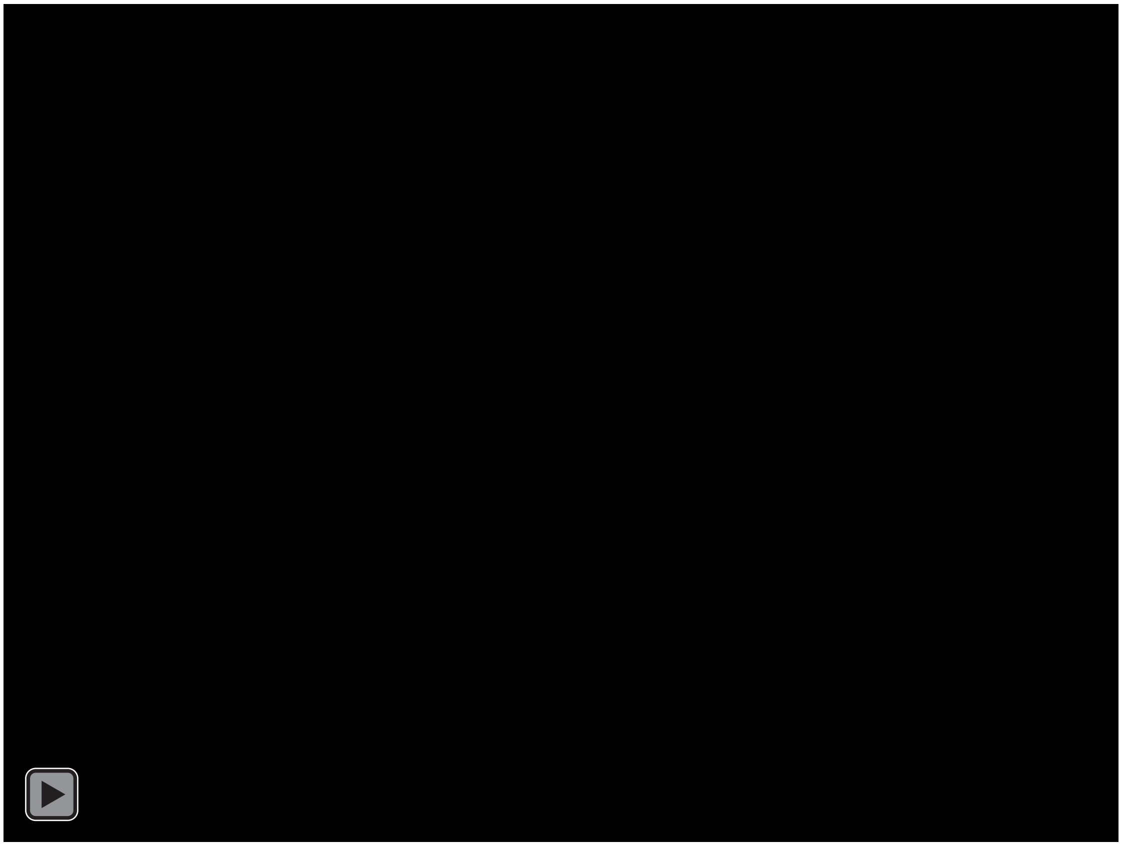

**Movie S2.**

yKD1177 (SMC1-AID) EthOH, cell repairs, resynthesises the red spot and divides

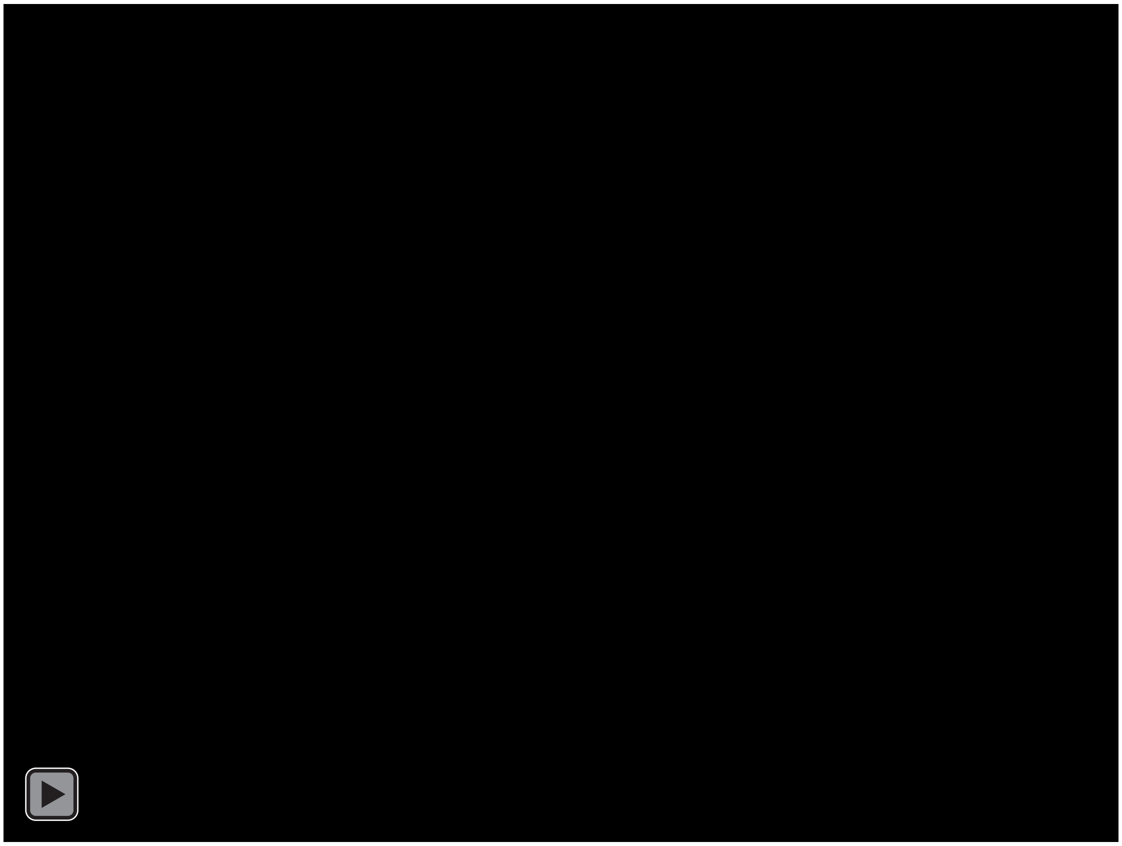

**Movie S3.**
yKD1177 (SMC1-AID) EthOH, no repair, no division

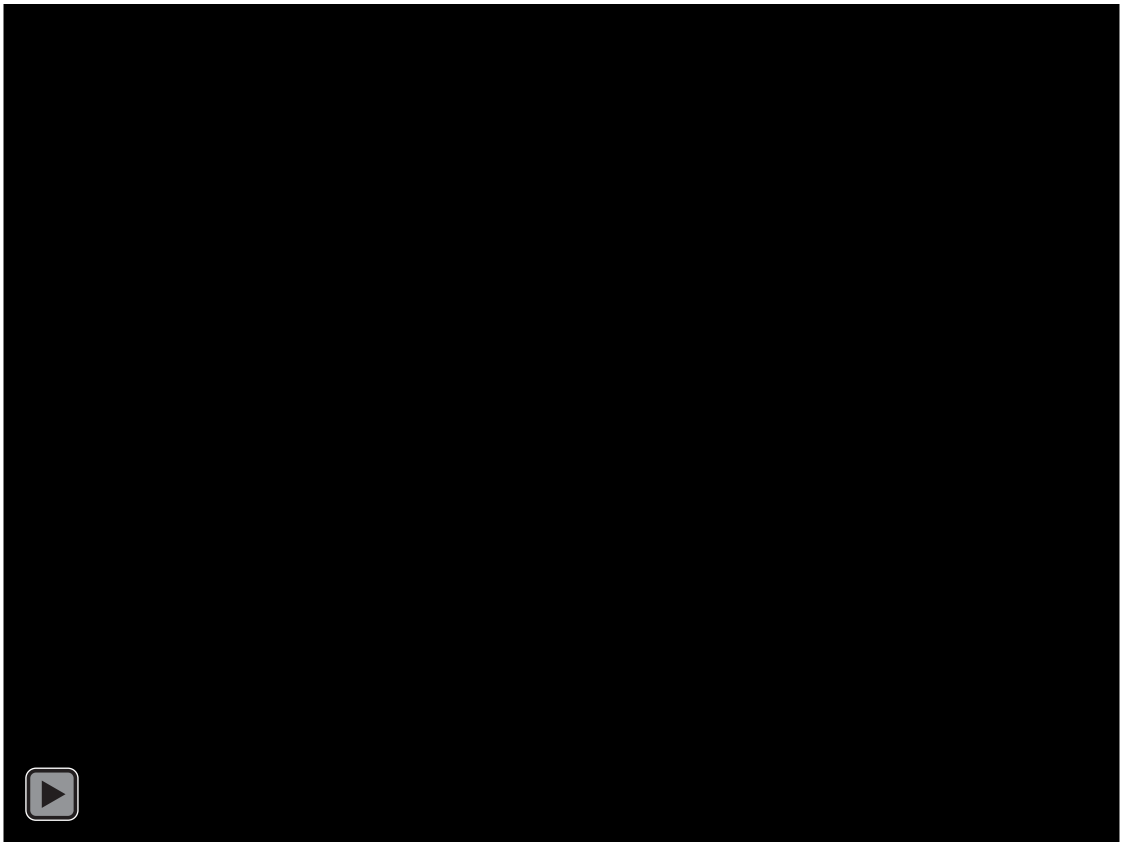

**Movie S4.**

yKD1177 (SMC1-AID) EthOH, no repair, division

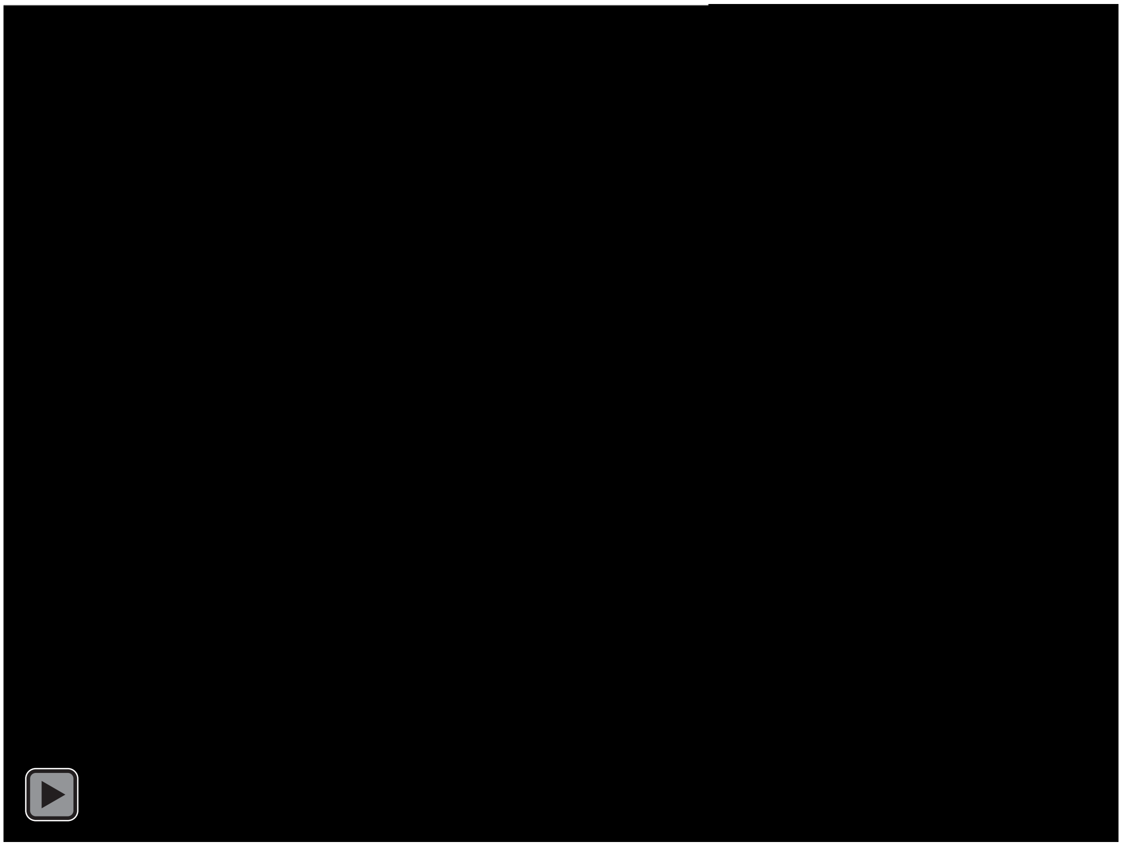
